# SiSaNA: A Python-based command line interface for Single-Sample Network Analysis

**DOI:** 10.1101/2025.11.06.680212

**Authors:** Nolan K. Newman, Tatiana Belova, Marieke L. Kuijjer

## Abstract

**Motivation:** Reconstructing and analyzing single-sample regulatory networks has proven useful for understanding mechanisms of regulatory biology and its disruption in disease, but traditionally required a fair amount of programming experience and work from the user.

**Results:** Here, we present SiSaNA (<u>S</u>ingle-<u>S</u>ample <u>N</u>etwork <u>A</u>nalysis), a workflow that removes the need of prior programming experience and allows for single-sample regulatory network reconstruction and analysis, all via a command line interface. Using example input files, we walk the user through a tutorial on how to run the SiSaNA pipeline, interpret the results, and obtain high-quality figures summarized in a single HTML file as output.

**Availability:** SiSaNA is freely available to install with pip via the Python Package Index. It is additionally available on GitHub at https://github.com/kuijjerlab/sisana. Example input files can be found at https://doi.org/10.5281/zenodo.17190642.

## Introduction

Reconstructing gene regulatory networks allows for the inference of transcription factor (TF) regulatory activity, the examination of gene targeting patterns, and individual regulatory interactions. Various tools exist to reconstruct and interrogate regulatory networks, including those in the Network Zoo (NetZoo) ecosystem(1). One tool included in NetZoo is PANDA(2) (Passing Attributes between Networks for Data Assimilation) which uses a message-passing technique to incorporate prior knowledge on putative gene regulation (whether TFs can bind to regulatory regions of genes) and TF-TF interactions with gene co-expression data to generate a single bipartite regulatory network for multiple samples. Source nodes in these networks are TFs and target nodes are genes, with edge weights representing the likelihood of regulation of TFs on target genes. LIONESS(3) (Linear Interpolation to Obtain Network Estimates for Single Samples) is another tool in the NetZoo ecosystem, which can be applied to the PANDA algorithm to reconstruct single-sample regulatory networks. Single-sample regulatory networks modeled with PANDA and LIONESS have proven useful for studying various aspects of biology, including e.g., identification of sex-specific regulatory differences in colon cancer(4), regulatory alterations associated with brain cancer survival(5), and a therapeutic candidate for Merkel cell carcinoma(6).

Despite their clear benefits, using single-sample regulatory network reconstruction algorithms can be challenging as they require a decent amount of computational proficiency, both for the reconstruction and downstream analysis of the single-sample networks. Therefore, we have developed a new command line interface tool, SiSaNA (<u>S</u>ingle-<u>S</u>ample <u>N</u>etwork <u>A</u>nalysis) that seamlessly integrates into the NetZoo ecosystem. SiSaNA utilizes the PANDA and LIONESS algorithms from NetZoo, but provides a more user-friendly experience for reconstructing, comparing and interpreting single-sample regulatory networks, all via the command line. SiSaNA requires little to no programming proficiency other than basic familiarity with the command line, while providing high-quality figures as output.

## Overview of the SiSaNA pipeline

The SiSaNA GitHub repository outlines the installation steps required for installing from the Python Package Index (PyPI). After installing from PyPI, one can run the **sisana -e** command to download example input files from the SiSaNA Zenodo repository. An example analysis using these files follows below.

SiSaNA works in multiple steps, each of which have been laid out in **Figure 1**. These are also described on the GitHub page and SiSaNA command line help documentation. Each step is labeled with a command (such as “preprocess” or “generate”).

**Fig. 1.**
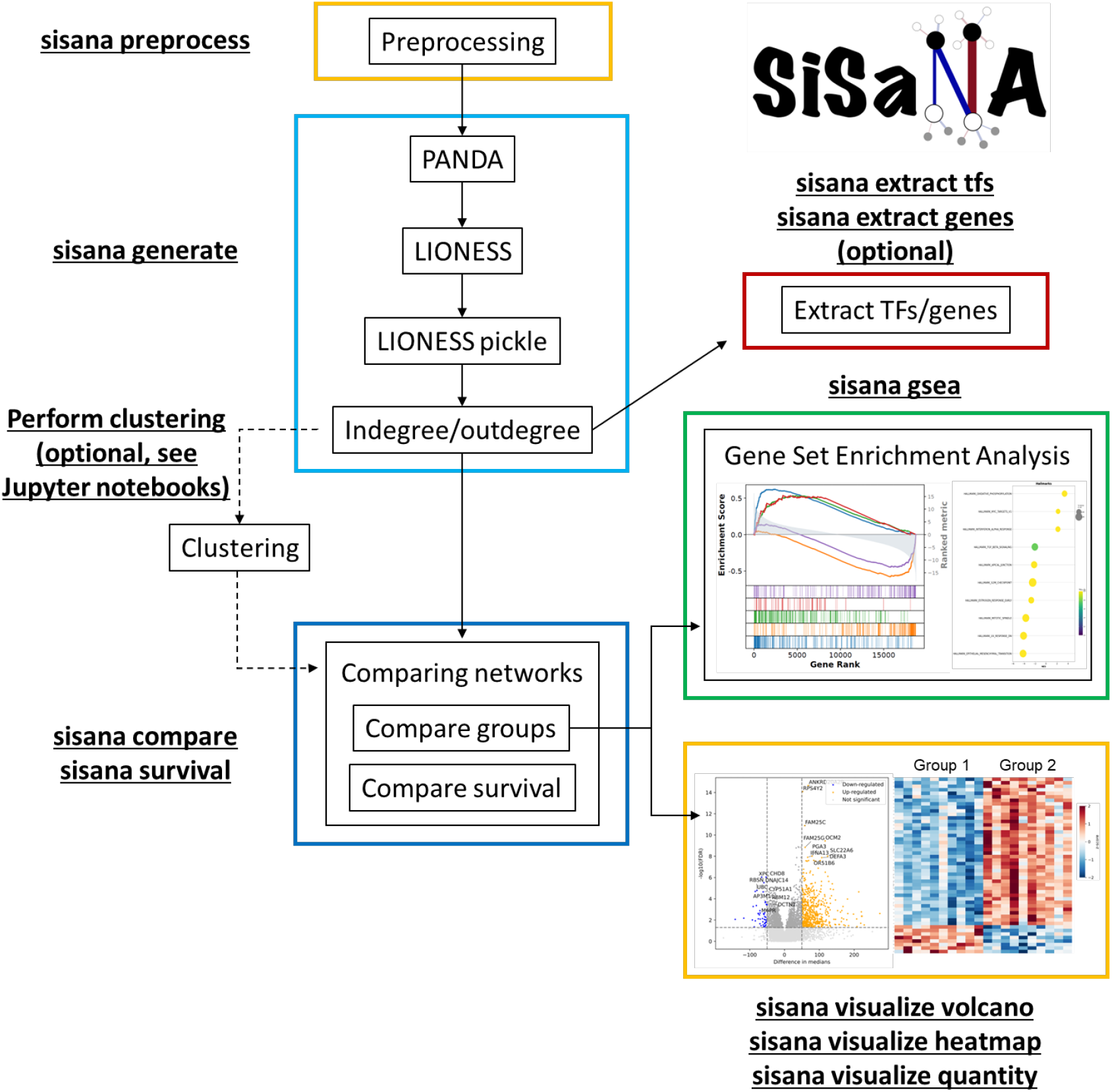
Overview of the SiSaNA pipeline. SiSaNA contains multiple commands, including “preprocess”, “generate”, etc. These commands are used to tell SiSaNA which section of the pipeline the user wishes to run. Some commands (e.g. “extract” and “visualize”) have additional subcommands that need to be specified to tell SiSaNA the specific type of comparison or visualization to perform. For example, the command “sisana visualize volcano” will create a volcano plot of the data, using the parameters set in the params.yml file. More information and examples of how to set up the params.yml file of SiSaNA can be found on the GitHub page.

### Preprocessing of input data using SiSaNA

The **preprocess** command requires three files as input: 1) pre-normalized, log-transformed transcriptomic data in standard matrix form (samples as columns, genes as rows), 2) a protein cooperativity file including putative protein-protein interactions (PPI) between transcription factors, and 3) a prior motif file denoting whether there is a putative transcription factor binding site in the promoter region of a gene. SiSaNA requires the naming format of TFs and genes to be consistent across all three files. The PPI and prior motif files can be generated using e.g. SPONGE(7). However, pre-generated PPI and motif files for Homo sapiens can be found in the SiSaNA Zenodo repository. The input files/parameters for this step and all future steps are defined in the params.yml file (see GitHub page and provided example parms.yml file for more information and specific examples).

### Using SiSaNA to reconstruct single-sample networks

The **generate** command utilizes the netZooPy package(1; 3; 8) (part of the Network Zoo(1) ecosystem of tools for gene regulatory network analysis) to reconstruct models using PANDA and LIONESS. The PANDA algorithm uses a message-passing technique to reconstruct an aggregate regulatory network from multiple samples. When combined with PANDA, the LIONESS algorithm(3) reconstructs a network using all samples (*n*). It then iteratively removes single samples (*q*) from the sample pool one at a time (*n*-*q*), and performs the PANDA method after each sample removal. The edge score for sample *q* is calculated by subtracting the edge scores from the network *n*-*q* from the original aggregate network *n*, multiplying this by a scaling factor, and then adding these values to the edge scores from the network modeled on *n*-*q*. SiSaNA automatically serializes the output of the single-sample networks to a.pickle file, allowing for faster loading of the large file generated by the approach in downstream steps.

In the output file, edges are weighted in the resulting single-sample networks and can be interpreted as the likelihood (larger edge value equals greater likelihood) of regulation of a target gene by a TF. After all single-sample networks have been modeled, gene in-degrees (also known as gene targeting scores(9)) and TF out-degrees will be calculated. Importantly, as networks modeled with PANDA are complete and weighted, the in-degree values are based on summing up edges from all TFs to a specific gene, while the out-degrees sum up edges from a TF to all genes. A high in-degree means that a gene has a high likelihood of being regulated. However, the value does not indicate whether the gene is being activated or repressed. A high out-degree corresponds to a higher overall likelihood of regulation, which could mean a transcription factor targets many different genes, or potentially regulates only a few genes, but to a great extent.

### Extracting specific edges from the network

To determine the likelihood to which a single TF regulates its target genes (based on the edge weights connected to that TF), the **extract tfs** command-subcommand can optionally be used. This will extract the per-sample edge weights of the specified TFs and their target genes, then save the resulting edge weights in a text file. This is useful if the user is interested in exploring the regulation of specific genes or the amount of targeting by specific TFs. Likewise, the **extract genes** command-subcommand can be used to extract all edges that connect to specific genes.

### Downstream analysis of single-sample regulatory networks

If sample groups are predefined in the input data (e.g. “Control” and “Disease” samples), users can continue to the next step. However, if undefined, users may wish to employ various clustering algorithms to identify sub-groups of samples based on network properties (e.g. in-degree). As this step relies heavily on user interaction with the dataset, this analysis is more easily performed in an alternative environment, such as a Jupyter notebook. As such, we have supplied example Jupyter notebooks on GitHub (https://github.com/kuijjerlab/sisana/blob/main/clusteringtemplate.ipynb) and in the example files on Zenodo that aid in identifying subgroups via various clustering methods, including hierarchical clustering, k-means, and PCA/tSNE plots.

Once users have defined sample groups for comparative analysis, either predefined through known sample characteristics, through the use of unsupervised clustering algorithms, or identified through other techniques such as dimensionality reduction, they can continue to the **compare** command. This command compares the expression or degrees of specific TFs/genes in these groups statistically with parametric or non-parametric tests, i.e. using a Student’s t-test or Mann-Whitney test for unpaired, or paired t-test or Wilcoxon signed-rank test for paired comparisons. Users can specify the type of test to perform in the params.yml file. A survival analysis can also be performed using SiSaNA with the **survival** command, allowing for users to test whether their sample groups differ in survival times (e.g. for users studying disease outcome).

For visualization of differences in e.g. in-degree, out-degree, or expression between groups, the **visualize** command can be used. Using this command, the user can create volcano plots (**visualize volcano**), heatmaps (**visualize heatmap**)(**Figure 2A-B**), box plots, and violin plots (**visualize quantity**), allowing for a simple way to contrast different sample groups.

**Fig. 2.**
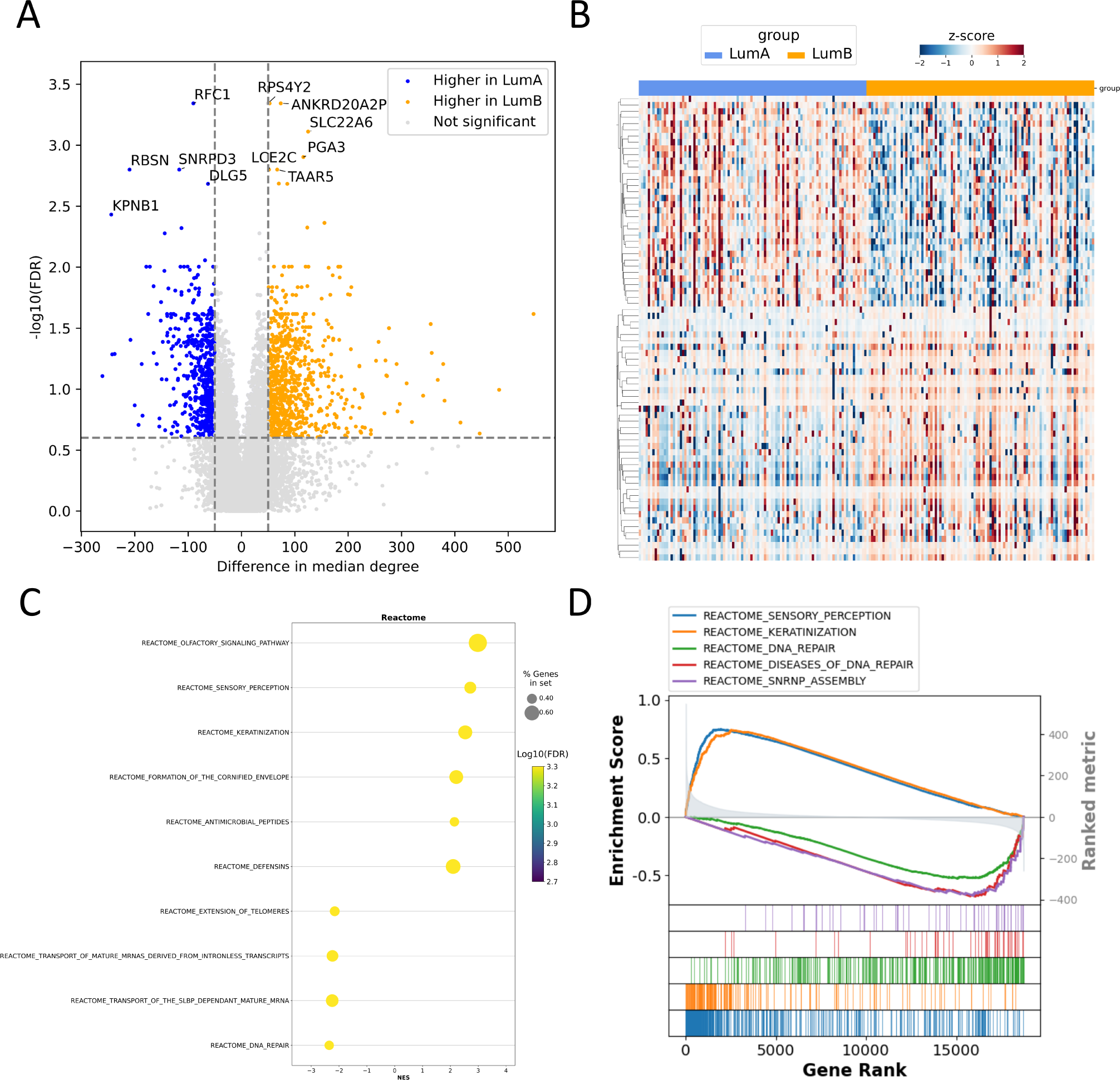
Example outputs of the comparison between LumA and LumB breast cancer samples, produced by SiSaNA. **A)** A volcano plot comparing the in-degree of genes between the LumA and LumB groups. Labeled genes are the top differentially regulated genes based on a combination of the difference in median degree and −log10(FDR) **B and C)** A gene set enrichment analysis(10)(11) (from the GSEApy package, which is incorporated in the SiSaNA pipeline) of the top differentially regulated genes (based on in-degree) between the LumA and LumB groups to identify enriched pathways those genes are involved in. These are the pathways that are differentially regulated between the two groups. **D)** A clustermap, clustered on genes, that is used for visualizing the top genes (based on in-degree) that differentiate the two groups.

To explore these groups further, one can perform a gene set enrichment analysis(10) in SiSaNA by using the **gsea** command. SiSaNA uses the GSEApy package(11) in combination with the statistics calculated in the previous **compare** command to perform the gene set enrichment analysis, then outputs the GSEA enrichment plot and dot plot (**Figure 2C-D**), along with other files created by GSEApy. GSEApy uses files in GMT (Gene Matrix Transposed) file format. Example GMT files (KEGG MEDICUS subset of CP, Reactome subset of CP, and Hallmark) from the human MSigDB collection can be downloaded together with the other example files described above using the **sisana -e** command, but users can also supply other GMT files if desired.

The analysis and all resulting plots from each of these steps are summarized in an HTML file (Supplementary File 1).

## Using SiSaNA to compare subtypes in breast cancer: an example analysis workflow

Breast cancer is a heterogeneous disease, consisting of multiple defined subtypes, commonly classified from the PAM50 gene signature(12). With this gene signature, tumors are classified into one of five distinct subtypes: basal-like, HER2-enriched, normal-like, luminal A, and luminal B breast cancer. Here, we present an example of using SiSaNA, in which we will compare single-sample network characteristics between the luminal A (LumA) and luminal B (LumB) samples from The Cancer Genome Atlas (TCGA) breast cancer (BRCA) cohort. We will then identify enriched biological pathways based on the regulation of genes (in-degree).

### Example steps for running SiSaNA

Example input files can be obtained by running the command **sisana -e**, which downloads the example files from the SiSaNA Zenodo repository into a new directory called **example inputs**. For an exhaustive list of all parameters and how to use them, please see the example params.yml provided in this directory. Example commands for each step can be found in the help documentation of each command by using the **-h** flag. An explanation of each step for this example analysis of generating and comparing single-sample regulatory networks is given below, including the parameters used in the params.yml file:

1. **sisana preprocess./example inputs/params.yml** First, filter out genes not expressed in a sufficient number of samples. In this example, we will use five (out of the 20 input samples) for the “number” parameter to indicate the number of samples that a gene must be expressed in. Filtering is important, as network reconstruction with PANDA relies on co-expression measurements, and each gene must show some variation in expression across samples.
2. **sisana generate./example inputs/params.yml** Next, use the PANDA and LIONESS algorithms (specified with “lioness” for the “method” parameter) to reconstruct networks. Then, calculate the gene in-degrees and TF out-degrees. Specifically, we will use “cpu” (instead of “gpu”) for the compute parameter, and use 20 for the number of cores (ncores parameter). Users will need to modify this value depending on the hardware being used. By default, the “start” and “end” parameters are left blank. These options are only needed if one wishes to reconstruct networks for a subset of samples, for example, to analyze a subset modeled against a larger background, or to generate multiple subsets of networks in separate runs, which can be combined at a later stage with the **combine** command. Leaving these parameters blank will reconstruct networks for all samples.
3. **sisana compare./example inputs/params.yml** Here, we compare the in-degrees and out-degrees (by using “degree” for the “datatype” parameter) for the LumA vs LumB samples (specified in the “groups” parameter) to find the top differentially regulated genes. We specify we want to use the Mann-Whitney U test (instead of a Student’s t-test) by setting “mw” for the “testtype” parameter. The.rnk file will be sorted by the median difference between the two groups, as specified by using “mediandiff” for the “rankby” parameter.
4. **sisana gsea./example inputs/params.yml** Next, we will perform a gene set enrichment analysis between the sample groups to identify pathways that distinguish each group. Here, we have specified the GMT file to use (“Hallmark.v2023.2.Hs.symbols.gmt”) in the “gmtfile” parameter and indicated to SiSaNA that we wish to call this gene set “Hallmarks” (in the “geneset” parameter), which is used for labeling the output file name.
5. **sisana visualize volcano./example inputs/params.yml** Finally, we will visualize differences in in-degree between the LumA and LumB groups with a volcano plot. Here, we specify that we want to plot the difference in median degree (“difference_of_medians_(LumA-LumB)”) on the x-axis (with the “diffcol” parameter) and use the “FDR” column as the “adjpcol” parameter. We then must also specify a threshold for coloring genes in each group on the x-axis (with the “xaxisthreshold” parameter) and y-axis (with the “adjpvalthreshold” parameter). We have set 50 (which corresponds to *<* (−50) or *>* 50) and 0.25 as our thresholds respectively for this example. Finally, in this example we assumed the user had some set genes they were interested in labeling, so we gave a list of genes to the “genelist” parameter. If we had not done that, then we would have needed to set the “numlabels” parameter instead, which dictates how many genes on each side of the volcano plot to label.
6. **sisana summarize** Summarize the outputs of SiSaNA into a single HTML file. Please note that this command must be ran from the directory containing the log files subdirectory.

### Results of the example analysis

Despite the similarities between LumA and LumB breast cancer, we are still able to identify genes that are differentially regulated (meaning they have significantly different values of in-degree) between these breast cancer subtypes. Genes higher in the LumA group include *RFC1, RBSN, SNRPD3, DLG5*, and *KPNB1*, among others. Meanwhile, genes more highly regulated in the LumB group include *RPS4Y2, ANKRD20A2P, SLC22A6, PGA3, TAAR5*, and *LCE2C*, among others (**Figure 2A**). Additionally, we can cluster the top differentially regulated genes to identify those that best distinguish LumA patients from LumB patients (**Figure 2B**).

Finally, based on the gene set enrichment analysis of the in-degree values, we can identify various enriched pathways specific to the LumA group (keratinization, sensory perception, and formation of the cornified envelope) and the LumB group (mitotic prometaphase and cell cycle) (**Figure 2C-D**). These differences in regulatory network structure highlight subtype-specific transcriptional programs, with LumA enriched for pathways linked to epithelial differentiation and LumB showing stronger regulation of cell-cycle-related processes. For further downstream analysis, a user could use the **sisana extract** command to extract just those genes (and their associated edges to transcription factors) and compare edge weights between sample groups.

## Example data and analysis methods

Pre-generated PPI and motif files for Homo sapiens were downloaded from the SPONGE Zenodo repository (https://doi.org/10.5281/zenodo.15063580)(7).

Example expression input data was downloaded and preprocessed as described in Belova et al.(13). To make the example analysis computationally lightweight and suitable for demonstration purposes, we selected a subset of the data for input to the **preprocess** and **generate** steps. Specifically, we filtered the expression data to only include 5,000 genes. Then, we selected ten random LumA samples and ten random LumB samples to include as input to the **generate** step.

In consideration for having sufficient statistical power for all SiSaNA steps downstream of **generate**, we used the indegrees from 100 LumA and 100 LumB individuals (randomly selected) that were calculated in Belova et al.(13). The exact commands and parameters used are described in the section **Example steps for running SiSaNA** and in the example params.yml file.

## SiSaNA performance

To determine an estimated amount of time SiSaNA requires to run, we timed how long it took to run the example files from Zenodo through the pipeline. A machine running the Red Hat Enterprise Linux distribution with an AMD EPYC 7643 48-Core processor and 2 TB of RAM was used. The “date” command was used before and after the pipeline and the difference in times was then used as the amount of time the pipeline took. The performance was evaluated using the example input dataset, which includes 20 samples for the **preprocess** and **generate** step, then 200 samples for the remaining steps. To run the steps **preprocess, generate, compare, survival, gsea, visualize volcano**, and **visualize heatmap** it took six minutes and seven seconds in total. Unsurprisingly, the longest step was generating the networks, which took four minutes and 37 seconds when running on the CPU.

## Conclusion

SiSaNA is a new computational tool that interfaces with netZooPy, allowing for preprocessing of data prior to running single-sample gene regulatory network modeling with PANDA and LIONESS and downstream analysis of the single-sample regulatory networks. It provides a number of features and benefits for comparing and reconstructing regulatory networks:

- SiSaNA is easy to obtain and use, requiring **little to no programming experience**, only basic familiarity with the command line on UNIX-based systems.
- SiSaNA is written completely in Python and distributed with PyPI, allowing for ease of installation and management in a conda environment.
- Basic network properties such as in-degrees (also known as gene targeting scores) and out-degrees can be calculated.
- With SiSaNA, one can easily compare and visualize network properties to find genes that are differentially regulated between sample groups, perform survival analyses between groups, and/or identify enriched sets of genes.

## Limitations

As SiSaNA is a command line based tool, the options for users are more limited than they would be if users were to write their own code and work interactively with PANDA and LIONESS (or other tools for reconstructing gene regulatory networks). Additionally, although we have tried our best to remove any requirements for users to be familiar with programming, users without pre-defined sample groups may need to do some work in Jupyter notebooks, although we have tried to mitigate the complexity of that work by providing example Jupyter notebooks users can use as reference.

## Author contributions statement

N.N. developed the SiSaNA software and wrote the manuscript.

T.B. and M.K. contributed ideas for the software and reviewed the manuscript.

## Acknowledgments

This work was supported by funding from the Research Council of Norway, Helse Sør-Øst, and the University of Oslo through the Norwegian Centre for Molecular Biosciences and Medicine (NCMBM) [187615], the Research Council of Norway [313932], the Norwegian Cancer Society [214871, 273592], the *‘Familien Blix’ fond til fremme av medisinsk forskning*, and the iCAN Flagship in Digital Precision Cancer Medicine.

We would like to thank Ine Bonthuis and Iga Niemiec for testing and proof-reading the SiSaNA tool and manuscript, as well as Romana Pop for testing the code.

## References

1. Ben Guebila, M., et al., The Network Zoo: a multilingual package for the inference and analysis of gene regulatory networks. Genome Biology, 2023. 24(1): p. 45.

2. Glass, K., et al., Passing messages between biological networks to refine predicted interactions. PloS one, 2013. 8(5): p. e64832.

3. Kuijjer, M.L., et al., Estimating sample-specific regulatory networks. Iscience, 2019. 14: p. 226–240.

4. Lopes-Ramos, C.M., et al., Sex differences in gene expression and regulatory networks across 29 human tissues. Cell reports, 2020. 31(12).

5. Lopes-Ramos, C.M., et al., Regulatory network of PD1 signaling is associated with prognosis in glioblastoma multiforme. Cancer research, 2021. 81(21): p. 5401–5412.

6. Yang, J., et al., Integrative analysis reveals therapeutic potential of pyrvinium pamoate in Merkel cell carcinoma. bioRxiv, 2023.

7. Hovan, L. & Kuijjer, M. L. (2025). SPONGE: Simple Prior Omics Network GEnerator. Bioinformatics, 2025, btaf320.

8. van IJzendoorn, D.G., et al., PyPanda: a Python package for gene regulatory network reconstruction. Bioinformatics, 2016. 32(21): p. 3363-3365.

9. Weighill, D., Ben Guebila, M., Glass, K., Platig, J., Yeh, J. J., & Quackenbush, J. (2021). Gene targeting in disease networks. Frontiers in Genetics, 12, 649942.

10. Subramanian, A., et al., Gene set enrichment analysis: a knowledge-based approach for interpreting genome-wide expression profiles. Proceedings of the National Academy of Sciences, 2005. 102(43): p. 15545–15550.

11. Fang, Z., X. Liu, and G. Peltz, GSEApy: a comprehensive package for performing gene set enrichment analysis in Python. Bioinformatics, 2023. 39(1): p. btac757.

12. Parker, J. S., Mullins, M., Cheang, M. C., Leung, S., Voduc, D., Vickery, T.,…& Bernard, P. S. (2009). Supervised risk predictor of breast cancer based on intrinsic subtypes. Journal of clinical oncology, 27(8), 1160–1167.

13. Belova, T., Hsieh, P. H., Hovan, L., Osorio, D., Kuijjer, M. L. (2025). Pan-cancer analysis of patient-specific gene regulatory landscapes identifies recurrent PD-1 pathway dysregulation linked to outcome. bioRxiv, 2025–10.

