## Supplementary File 1 for "SiSaNA: A Python-based command line interface for Single-Sample Network Analysis": results_summarized.html

Python Results


### SiSaNA results

#### Preprocessing

After removing **93** genes that were not present in at least 5 samples, **4908** genes were retained for the network reconstruction step.

#### Survival plot

Your groups differ in survival with a p-value of **0.330**. This indicates that there is a **non-significant** difference in the survival between the groups.

#### Volcano plot

To interpret this plot, pay attention to the TFs/genes that are colored. These are the genes that are below the **0.25** FDR threshold and greater (absolute value) than the threshold of **50** set for the difference in median degree. These may be genes that are important in distinguishing your two groups from one another.

There are **580** higher in LumA and **823** genes higher in LumB.

#### Quantity plot

Below you will find the heatmap created for visualizing the genes. Column clustering was not performed. Row clustering was performed.

#### GSEA results

Below you will find the top pathways that are enriched between the two sample groups. The GMT file used for generating these results was c2.cp.reactome.v2023.2.Hs.symbols.gmt.
